# Structural and energetic analysis of stabilizing indel mutations

**DOI:** 10.1101/2024.12.18.629072

**Authors:** Yulia M. Gutierrez, Gabriel J. Rocklin

## Abstract

Amino acid insertions and deletions (indels) are among the most common protein mutations and necessitate changes to a protein’s backbone geometry. Examining how indels affect protein folding stability (and especially how indels can increase stability) can help reveal the role of backbone energetics on stability and introduce new protein engineering strategies. Tsuboyama et al. measured folding stability for 57,698 single amino acid insertion or deletion mutants in 405 small domains, and this analysis identified 103 stabilizing mutants (ΔΔG_unfolding_ > 1 kcal/mol). Here, we use computational modeling to analyze structural and energetic changes for these stabilizing indel mutants. We find that stabilizing indel mutations tend to have local structural effects and that stabilizing deletions (but less so insertions) are often found in regions of high backbone strain. We also find that stabilizing indels are typically correctly classified as stabilizing by the Rosetta energy function (which explicitly models backbone energetics), but not by an inverse folding (ESM-IF)-based analysis (Cagiada et al. 2024) which predicts absolute stability (ΔG_unfolding_).

## Introduction

Amino acid insertion and deletion (indel) mutations are among the most common sources of variation in proteins (*1*–*3*). Although the indel mutation rate varies by species and indel length (*4, 5*), indel mutations are frequently implicated in diseases, such as the CFTR delF508 variant in cystic fibrosis (*6*) and the APP delE693 variant in Alzheimer’s disease (*7*–*9*). Unlike missense mutations, indels necessitate a change to a protein’s backbone, often causing more dramatic structural and energetic effects. While indel mutations are widely considered highly deleterious to protein folding and function (*10*–*12*), indels can also improve a protein’s activity and access sequence, structure, and functional space outside of the range allowed by point mutations (*13*–*17*).

Despite the prevalence of indels and their potential to accelerate protein engineering, the biophysics of indels are rarely studied and remain challenging to predict. Large-scale experimental techniques of mutational effects such as deep mutational scanning (DMS) and multiplex assays of variant effect (MAVEs) have primarily focused on the systematic characterization of amino acid substitutions (*18, 19*), although some recent studies have included indels (*8, 10, 16, 20*–*29*). Smaller scale indel mutagenesis studies have shown that indels are typically highly deleterious (*10, 21, 28*) and are more tolerated in loops and termini compared to secondary structure elements such as α-helices and β-strands (*2, 10, 21, 26, 30*). Although these studies provide important insights regarding indels, the energetic effects remain challenging to predict, in part due to few examples of tolerated indel mutations (*31, 32*). In 2024, Topolska et al. found that only 1% of indel mutants improved protein abundance compared to the wild-type protein (*10*). Moreover, in 2013 Dagan et al. identified a six-residue deletion in human muscle acylphosphatase that stabilized the enzyme by 4.3 kcal/mol (*33*). While stabilizing indel mutations are exceedingly rare, understanding these mutations can reveal the role of backbone energetics in determining protein folding stability. This would allow us to predict biological variant effects and ultimately leverage indel mutagenesis for protein engineering.

In 2023, Tsuboyama et al. measured the folding stabilities for thousands of small protein domains using their novel cDNA proteolysis method, including measurements for 57,698 single deletion and Ala or Gly insertion mutants (*34*). From these large-scale data, we identified 103 highly stabilizing indel mutations (ΔΔG_unfolding_ > 1 kcal/mol) in both natural and *de novo* designed proteins and investigated the structural and energetic effects of these mutations using a combination of computational techniques. ColabFold modeling revealed that stabilizing indels had primarily local effects on protein structure, and Rosetta analysis indicated that stabilizing deletions, but less so insertions, are commonly found in regions of high backbone strain. We also found that the Rosetta energy function outperforms the Cagiada ESM-IF analysis (*35*) (which does not explicitly model backbone energetics) in successfully classifying these stabilizing mutants. Finally, we identify favorable indel mutations that are common throughout the WW domain and SH3 domain families and describe the energetic basis of these favorable mutations. Along with our findings, these 103 stabilizing mutants can serve as an informative dataset to evaluate computational methods at predicting mutant effects on folding stability.

## Results & Discussion

### We identified 103 stabilizing indels in both natural and designed domains

Tsuboyama et al. used cDNA display proteolysis to measure the free energy of unfolding (ΔG_unfolding_) for hundreds of small protein domains, along with point mutations, glycine insertions, alanine insertions, and deletions at every position. For example, in the chromodomain of human chromobox protein homolog 7 (PDB ID: 2K1B, a component of the PRC1-like complex involved in regulating development (*34, 36*)), deleting any of the five residues highlighted in Figure 1A results in a more stable fold. Despite this example of highly stabilizing indel mutations, we find that indel mutations are typically more deleterious to a protein’s free energy of unfolding compared to substitution mutations (Fig. 1B). This result is consistent with current literature (*10, 20*).

**Figure 1.**
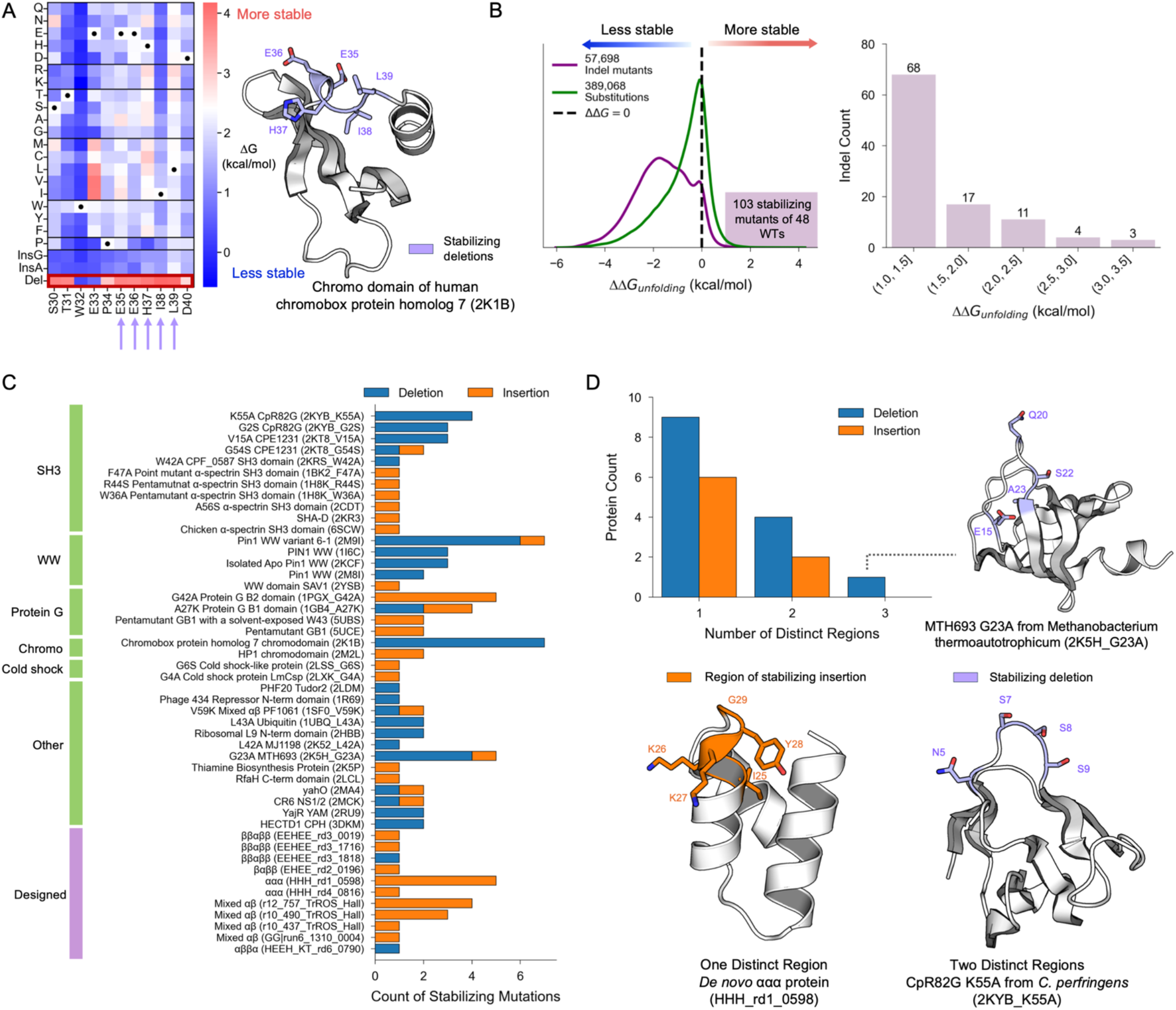
**A**, (left) Deep mutational scanning stability results of 2K1B from the Tsuboyama et al. Mega-scale dataset and (right) 2K1B structure highlighting residues that stabilize the protein upon deletion. Boxes are colored by ΔG_unfolding_, where red indicates a mutant more stable than the wild type and blue a mutant less stable. Five stabilizing deletions are highlighted with purple arrows on the heatmap, as well as on the wild-type 2K1B ColabFold model. **B**, (left) Distribution of ΔΔG_unfolding_ for all indel and substitution mutants. The purple box includes the 103 stabilizing indel mutants. (right) Distribution of ΔΔG_unfolding_ values for stabilizing indel mutants. **C**, All stabilizing indels organized by their wild-type domains. Insertions are colored in orange and deletions in blue. **D**, Distribution of the number of distinct regions in a wild-type protein that accommodate a stabilizing indel for wild types with more than one insertion or deletion. Distinct regions are illustrated through ColabFold models of 2K5H_G23A (three distinct regions), HHH_rd1_0598 (one distinct region), and 2KYB_K55A (two distinct regions).

From a set of 57,698 indel mutants with high-quality stability measurements (*34*), we identified 103 indels in 48 wild types that improved the wild-type protein’s global stability by at least 1 kcal/mol (Fig. 1B). Over 80% of stabilizing indel mutations improve a protein’s stability by less than 2 kcal/mol, and there are only three occurrences of indel mutations causing an increase in stability greater than 3 kcal/mol (Fig. 1B). Although the proteolysis data indicate that these mutations are all stabilizing, the magnitude of the effects may be underestimated owing to the limited dynamic range of the cDNA display proteolysis assay (approximately 0-5 kcal/mol) (*34*).

### Stabilizing indels occur in a diverse set of natural and wild-type domains

We identified stabilizing indel mutations across 48 wild-type proteins, including both naturally occurring and *de novo* designed small protein domains (Fig. 1C). More than 75% of wild-type proteins with at least one stabilizing indel mutation are natural domains, some of which have notable biological functions. One biologically relevant wild type is the CPH domain of human E3 ubiquitin ligase HECTD1 (3DKM) which is stabilized by deletions in two unique positions (Fig. 1 C, D). Overall, 25 wild-type domains are stabilized by more than one indel mutation. To understand the spatial relationship of stabilizing indels within one wild-type domain, we calculated the number of “distinct regions” in which stabilizing mutations were accommodated per wild type. We define a distinct region as an indel site with one or more residues separating the next mutation site of the same type (insertion or deletion). For proteins with more than one stabilizing indel mutation, these mutations tend to occur in adjacent regions in the protein. For example, the *de novo* triple helical bundle HHH_rd1_0598 is stabilized by consecutive insertions at positions 26 (A or G), 27 (A or G), and 28 (A only, G marginally stabilized the protein by 0.3 kcal/mol) (Fig. 1D). More than 60% of wild-type domains with multiple stabilizing deletions and 75% with stabilizing insertions have only one distinct region which accommodates stabilizing indels. One exception is the G23A mutant of MTH693 (PDB ID: 2K5H, PFAM: NfeD) of the anaerobic archaeon *M. thermoautotrophicum* (mutations were tested in the G23A background because the wild-type domain is too stable for the cDNA display proteolysis assay). This domain is stabilized by four deletion mutations in three distinct regions. However, all four mutations are found in the same extended loop connecting the first and second β-strands (Fig. 1D).

### Stabilizing indels cause minimal structural perturbations

We used predicted models to examine the structural trends of stabilizing indel mutations. Mutant and wild-type structures were predicted using ColabFold (*37*) and the top-ranking model was selected for analysis. Most mutants (92/103) had high confidence predicted structures (average pLDDT > 85) that were similar to the wild-type domain (average Cα RMSD > 5 Å). Among the remaining structures, eight had low confidence predictions and three showed large structural differences compared to the wild-type sequences. Because machine learning-based structure predictions are especially likely to fail on protein variants and *de novo* designed proteins lacking sequence alignments, we focused our further analysis on the 92 domains with confident predictions that are similar to the wild-type domain. Using the Dssp algorithm to assign secondary structural elements, we found that approximately 55% of stabilizing insertions and deletions occurred in loop regions of the protein (Fig. 2A). This is consistent with previous literature indicating higher indel tolerance in unstructured protein regions including loops and termini (*10, 21*). Surprisingly, we also found more than 30% of stabilizing indels occur in the middle of helices and strands (Fig. 2A).

**Figure 2.**
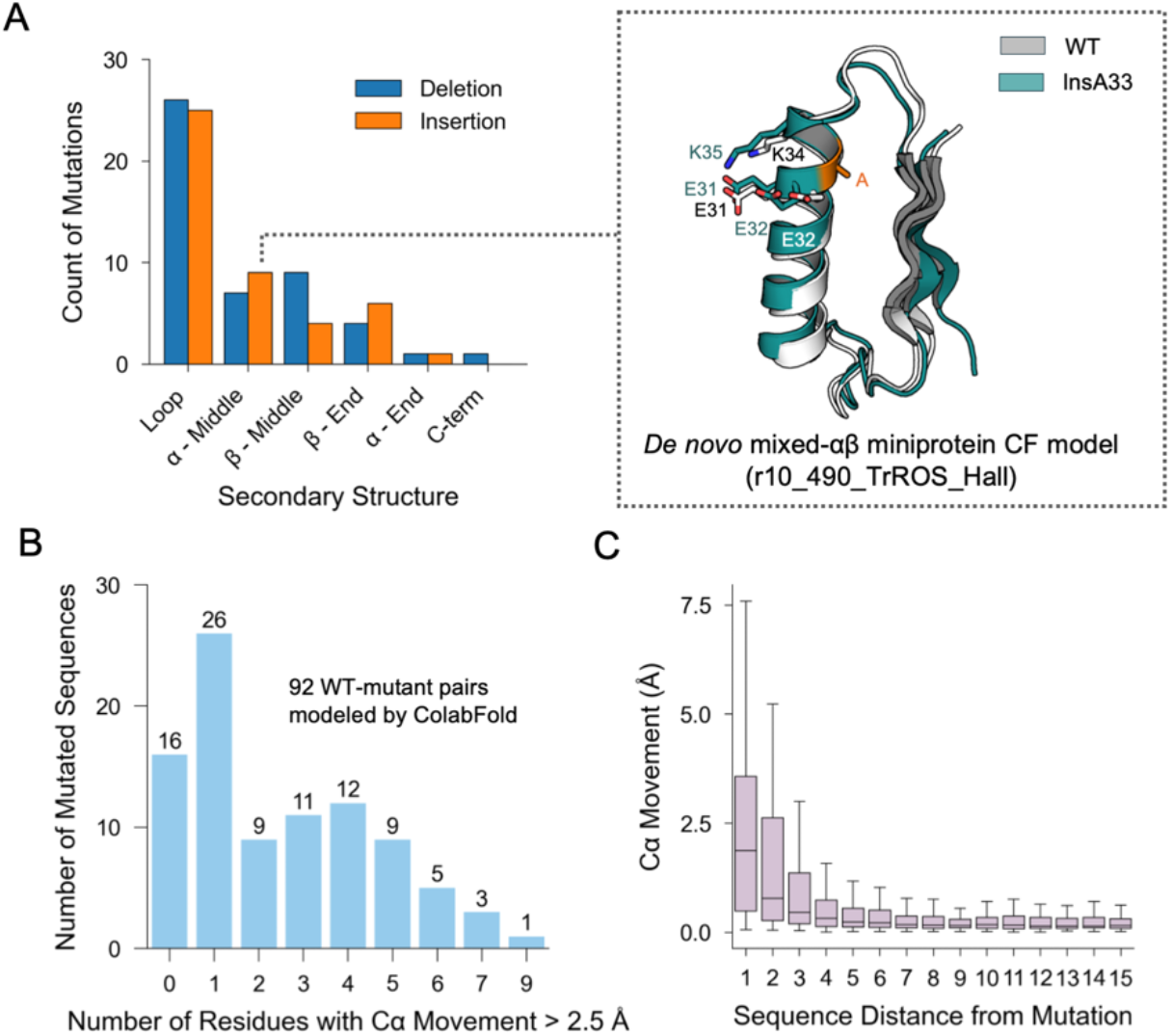
**A**, (left) Stabilizing indel mutations by secondary structure assignment. The end of an α-helix or β-strand is defined as directly adjacent to a loop residue, and the middle regions represent mutations not at the ends of an α-helix or β-strand. For deletions, the secondary structure assignments are DSSP classifications of the deleted residue, whereas insertions are the DSSP classifications of the inserted residue. (right) An example of an α-middle insertion is shown through ColabFold models of wild-type protein r10_490_TrROS_Hall (gray) and mutant InsA33 (teal). **B**, Distribution of the residues with Cα movement > 2.5 Å per mutated sequence for 92 wild type-mutant pairs modeled by ColabFold and aligned by LovoAlign. **C**, Distribution of Cα movement by separation in sequence from the site of indel mutation.

After removing wild type-mutant pairs with an average Cα RMSD greater than 5 Å, we characterized structural changes associated with stabilizing indel mutations. We find that stabilizing indels are predicted to cause minimal changes to the wild-type structure. We aligned ColabFold models of wild types and indel mutants using LovoAlign (*38*) and calculated the difference in each Cα position between the mutant and wild-type structures. More than 50% of wild-type-mutant pairs have two or fewer residues that changed position by more than 2.5Å, and approximately 90% had five or fewer residues that changed position by more than 2.5Å (Fig. 2B). Moreover, we observe that most large structural changes (Cα movement > 2.5Å) occur close in sequence to the stabilizing indel (Fig. 2C). In contrast, regions far from the mutation site typically are not predicted to undergo large structural changes (Fig. 2C).

### Rosetta outperforms ESM-IF in predicting mutational effects of stabilizing indels

We benchmarked two computational models, an ESM-IF-based analysis (*35*) and Rosetta (*39, 40*), at predicting the energetic effects of stabilizing indels. In 2024, Cagiada et. al developed an approach to predict stability (ΔG_unfolding_) based on sequence probabilities for a given backbone derived from the inverse folding model ESM-IF (*35*). First, we used the ESM-IF analysis to predict the ΔG_unfolding_ for each stabilizing indel mutant and its wild type. Then, we calculated the ΔΔG_unfolding_ for each mutant-wild type, where a positive value indicates the indel mutant is predicted to be stabilizing. By experimental measurements, all 92 indel mutants examined here are stabilizing, so positive ΔΔG_unfolding_ values are correct classifications. However, the ESM-IF analysis only predicts stabilizing effects for approximately half of the mutants (Fig. 3A). In the original Cagiada et al. model, ΔG_unfolding_ values are calculated as proportional to the sum of likelihoods for each amino acid in the analysis, so the predictions systematically increase with protein length. We also tested whether normalizing the predictions by sequence length would improve the predictions, but saw only a minimal effect (Fig. 3A).

**Figure 3.**
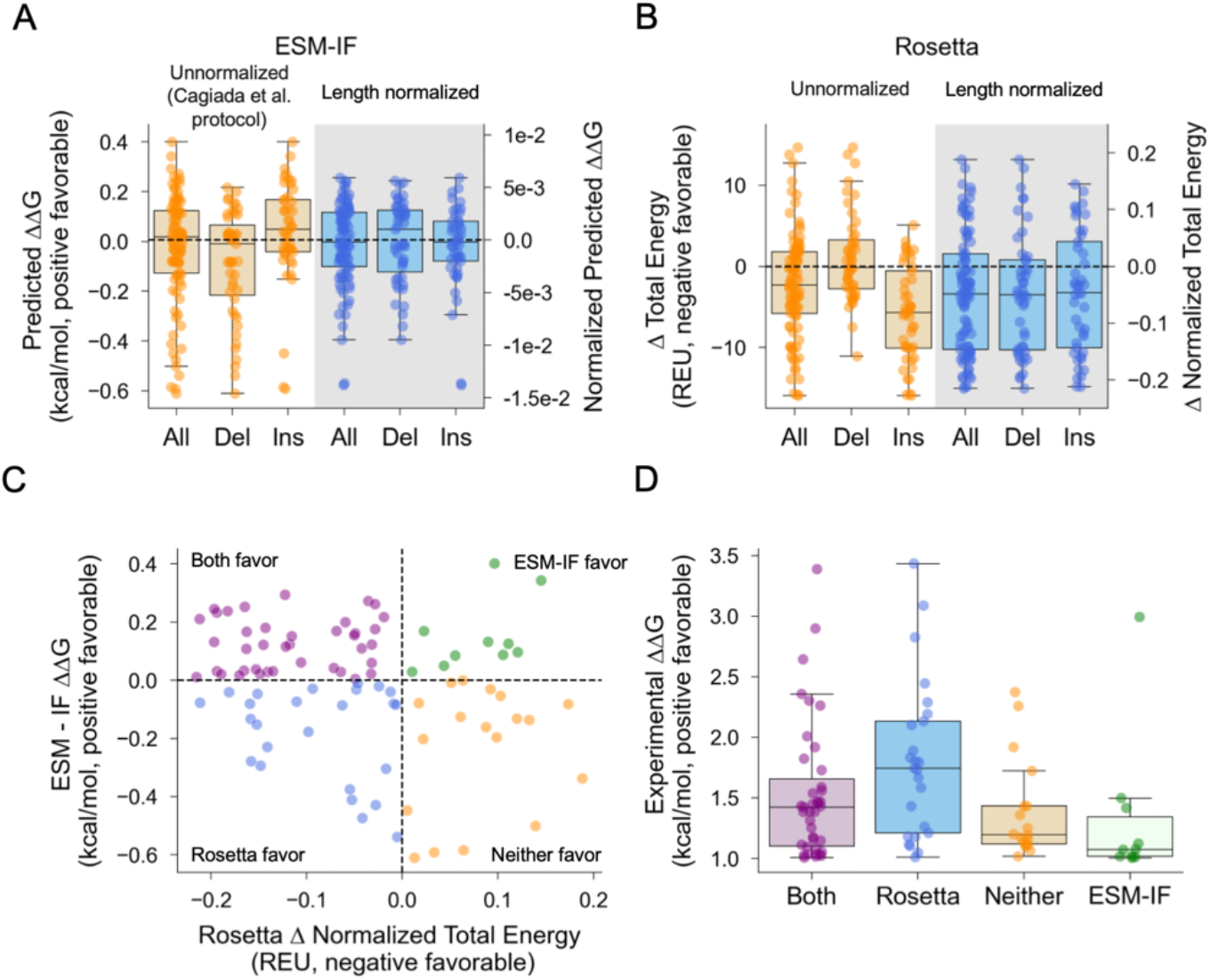
**A**, Distributions of unnormalized (left) and sequence length-normalized (right) ΔΔG_unfolding_ values predicted by the Cagiada ESM-IF analysis. Positive (favorable) ΔΔG values indicate accurate classifications. **B**, Distributions of unnormalized (left) and sequence length-normalized (right) Rosetta total score changes between indel mutant and wild type. Negative Δ Total Energy scores indicate accurate classifications. **C**, Relationship between ΔΔG_unfolding_ predicted by the Cagiada ESM-IF and the changes in normalized Rosetta total score. Points are colored by agreement of the Cagiada ESM-IF analysis and Rosetta. **D**, Distributions of experimental ΔΔG_unfolding_ for different categories of mutations based on Rosetta and Cagiada ESM-IF predictions.

We then used Rosetta, a physics- and knowledge-based model with a score function that explicitly calculates backbone energetics (*40*), to predict energetic changes for all 92 stabilizing indel mutants. We scored each protein with Rosetta and normalized total scores by protein length. For each predicted structure, we performed 20 relax runs using the Fast relax protocol. We calculated scores for each predicted structure by averaging individual score terms from the five lowest-scoring (by total energy) relaxed structures and normalized protein-level scores by dividing the score terms by protein length. Then, we calculated the difference in normalized total energy between the wild type and the indel mutant. A negative difference in normalized total energy indicates the indel mutant is predicted to be more favorable than its wild type. Rosetta predicts stabilizing effects for approximately two-thirds of mutants (Fig. 3B). Rosetta predicts the mutational effects of both insertion and deletion mutations equally well when total scores are normalized by sequence length (Fig. 3B).

Next, we investigated the agreement between Rosetta and ESM-IF predictions of changes in stability and their relationship to experimentally determined ΔΔG_unfolding_ values. Here, we compare ESM-IF ΔΔG_unfolding_ predictions to normalized Rosetta total scores because normalizing ESM-IF by sequence length had a minimal overall effect. Surprisingly, Rosetta and ESM-IF predictions showed only weak agreement (Pearson r = -0.23, Fig. 3C). We then divided all mutants into four categories according to whether they were predicted to be favorable by Rosetta, ESM-IF, neither, or both. We found that mutations favored by Rosetta alone were more experimentally stabilizing than mutations favored by both Rosetta and ESM-IF, and mutations favored by ESM-IF alone showed the lowest experimental ΔΔG values (Fig. 3C-D). Overall, these results highlight the moderate accuracy of Rosetta at modeling mutations that alter backbone structure, and the limitations of inverse folding models at capturing the effects of insertions and deletions.

### Stabilizing deletions (but not insertions) are often found in regions of backbone strain

We sought to understand the energetic effects of stabilizing indels using the Rosetta score function. Because indel mutations necessitate a change to a protein’s backbone, we hypothesized that indel mutations would occur in wild-type domains with a high degree of backbone strain. Using Rosetta, we scored 48 wild-type domains with at least one stabilizing indel mutation as well as 356 other wild-type domains in the Tsuboyama et al. dataset (*34*). We then calculated a backbone score for each protein, consisting of the sum of the “rama_prepro” and “p_aa_pp” score terms normalized to the length of the protein. The “rama_prepro” term describes the probability of a residue’s torsion angles Φ, Ψ given the amino acid identity while “p_aa_pp” refers to the probability of the amino acid identity given its torsion angles Φ, Ψ (*40*). Combined, these terms capture the overall favorability of a protein’s torsion angles, where residues with unfavorable angles represent regions of backbone strain. A higher backbone score suggests a higher degree of backbone strain. We find that wild-type proteins stabilized by indels do not have a higher degree of backbone strain than wild-type proteins not stabilized by indels (mean difference 0.0024 ± 0.027 (95CI computed by bootstrapping) Rosetta Energy Units (REU) per residue, Fig. 4A).

**Figure 4.**
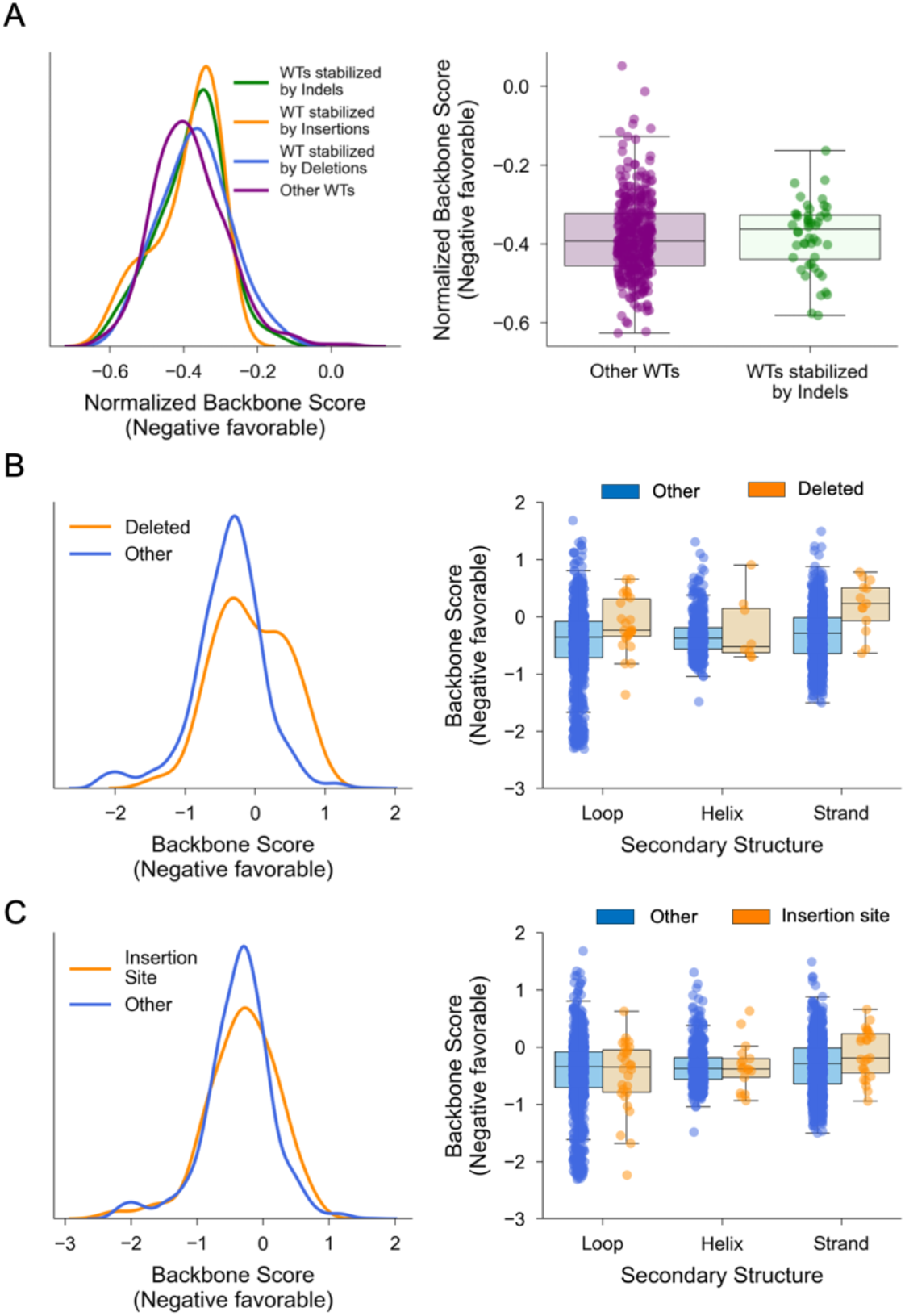
**A**, (left) Distributions of normalized Rosetta backbone scores for wild types stabilized by all indels (green), insertions (orange), deletions (blue), and all other wild-type proteins not stabilized by indels. (right) Comparison of normalized Rosetta backbone scores for wild types stabilized by indels and all other wild types. **B**, (left) Distributions of Rosetta backbone scores in the wild-type structure for deleted residues (orange) and other residues (blue) in 48 wild types stabilized by at least one indel mutation. (right) Rosetta backbone scores for deleted and other residues by secondary structure assignment. **C**, (left) Distributions of Rosetta backbone scores in the wild-type structure for sites of stabilizing insertions (the two residues flanking the insertion site, orange) and other residues (blue) in 48 wild types stabilized by at least one indel mutation. (right) Rosetta backbone scores for sites of stabilizing insertions and other residues by secondary structure assignment.

Although wild-type domains stabilized by indels were not considerably more strained than other wild-type domains, we suspected that stabilizing indels occur in regions of high local backbone strain. First, we investigated if sites of favorable deletions had high degrees of backbone strain in the wild type. Using our set of wild-type domains with at least one stabilizing indel, we calculated a per-residue backbone score (rama_prepro + p_aa_pp). We compared backbone scores at sites with favorable deletions to backbone scores at all other sites in the set of 48 wild-type domains with at least one stabilizing indel mutation. On average, deleted residues were 0.33 ± 0.14 (95CI) REU more strained than other residues. (Fig. 4B). Strand and loop residues showed the largest strain; favorably deleted strand residues were 0.50 ± 0.24 (95 CI) REU more strained than other strand residues, and favorably deleted loop residues were 0.34 ± 0.18 (95CI) REU more strained than other loop residues (Fig. 4B). However, favorably deleted helix residues showed nearly no difference in strain (average backbone score difference of 0.11 +/- 0.35 (95CI) REU).

Next, we investigated if stabilizing insertions occurred at strained sites in the wild-type protein. Here, we defined the stabilizing insertion site as the two residues in the wild-type protein that flank the inserted residue in the indel mutant. To understand if sites of stabilizing insertions are more strained than other protein sites, we compared the backbone scores of stabilizing insertion sites to all other sites in the 48 wild-type proteins with at least one stabilizing indel. On average, we find that sites of stabilizing insertions show no difference in strain compared to other sites (average backbone score difference of 0.0024 ± 0.18 (95 CI) REU) (Fig. 4C). Although stabilizing insertion sites are not more strained than other sites, we find that stabilizing insertions tend to occur in strained strands. On average, strand sites with stabilizing insertions were 0.20 ± 0.16 (95CI) REU more strained than other strand sites. In contrast, helix and loop sites with stabilizing insertions were not more strained than other sites, with average backbone score differences of 0.0045 ± 0.19 (95CI) REU and -0.020 ± 0.24 (95CI) REU, respectively (Fig. 4C).

### Pin1 WW domain is stabilized by loop deletions

The Pin1 WW domain illustrates how deletions can stabilize a domain by relieving backbone strain. Pin1 is a peptidyl-prolyl isomerase that isomerizes the phosphorylated Serine/Threonine-Proline (pSer/Thr-Pro) motif and is involved in numerous essential cellular processes, including cell division and differentiation (*41*). The WW domain of Pin1 is frequently used as a model system in protein folding studies (*42*–*45*), including in early mutational scanning experiments aimed at understanding stability (*46*). Our analysis identified deletions in Loop 1 of the Pin1 WW domain that are consistently stabilizing across four wild-type background variants (Fig. 5A, B). This loop plays a functional role in the WW domain by binding to a phosphorylated substrate (*47*– *49*), which may explain the selection for the wild-type sequence over the stabilized deletion variants. Notably, one of the variants identified by cDNA display proteolysis (delR12) was previously purified and found to increase the stability of the Pin1 WW domain by 1.0 kcal/mol (*42*). This supports the reliability of the cDNA display proteolysis data, which measured a 1.2 kcal/mol increase in stability.

**Figure 5.**
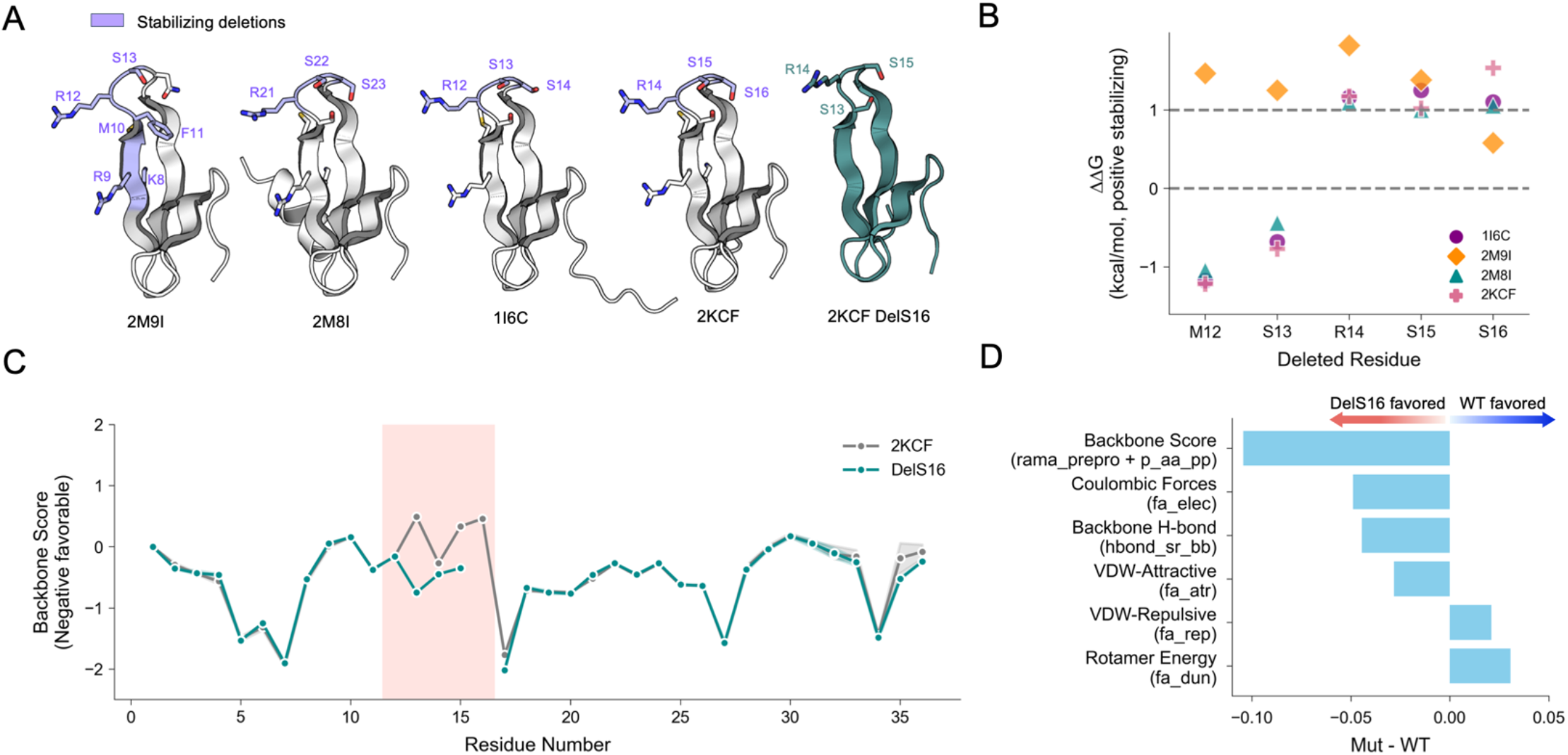
**A**, ColabFold models of four Pin1 WW domain wild-type background variants 2M9I, 2M8I, 1I6C, and 2KCF, as well as deletion mutant DelS16 2KCF. **B**, Experimental ΔΔG_unfolding_values for five corresponding deleted residues in 2M9I, 2M8I, 1I6C, and 2KCF. Residue numbering in 2KCF is used as a reference. **C**, Per-residue backbone score profiles for 2KCF and deletion mutant DelS16 2KCF. Residues 12-16 of 2KCF are highlighted in pink. **D**, Normalized changes in Rosetta score terms between deletion mutant 2KCF DelS16 and wild-type 2KCF.

To understand why deletions in Loop 1 stabilize this domain, we analyzed the individual energy terms of the Rosetta energy function. As before, we examined the backbone energy at each position and found that the stabilizing deletions are located in the most strained region of the wild-type domain. For example, S16 has the second-highest backbone score of all residues in the Pin1 WW domain (Fig. 5C). In the 2KCF_delS16 mutant, we observe a decrease in backbone score in neighboring residues S15 and S13, suggesting that deleting S16 both eliminates a strained residue and alleviates strain in nearby loop residues (Fig. 5C). On the domain level, Rosetta predicts the deletion mutant to be more favorable than the wild type, and the driving contributors are decreased backbone strain and increased electrostatic interactions (Fig. 5D). There are also smaller energetic contributions from changes in hydrogen bonding, Van der Waals forces, and side chain conformations (Fig. 5D). A previous analysis of mutations in Loop 1 noted that the wild-type loop structure (a type-II four-residue turn) is relatively rare, and that deletions can lead to the formation of a more common type-I G1 bulge turn, which is the main Loop 1 conformation in the WW domain family (*42*)

### Gly and Ala insertions can have different effects on protein stability

To better understand how Ala and Gly insertions differentially stabilize domains, we analyzed the effects of inserting either residue at 36 sites stabilized by an insertion. We found that Gly is nearly always tolerated at sites that are stabilized by an Ala insertion (ΔΔG > 0 kcal/mol), but Ala is often destabilizing at sites stabilized by a Gly insertion (Fig. 6A). The greater favorability and tolerability of Gly may originate from its greater conformational flexibility. At sites with stabilizing Ala insertions, ColabFold’s predicted structures typically model inserted Ala and Gly residue in the same region of the Ramachandran map (16/18 sites in the same ABEGO bin, Fig. 6A) (*50*). However, at sites stabilized by Gly insertions where Ala insertions do not meet our stabilizing threshold, ColabFold typically models Ala and Gly insertions to fall in different regions of the Ramachandran map (11/18 sites in different ABEGO bins, Fig. 6A). The group of sites stabilized by Ala insertions has a significantly higher proportion of ABEGO agreement between predicted Ala and Gly insertion conformations compared to sites only stabilized by Gly (Fisher’s exact test p = 0.0045). For sites stabilized by Gly insertions where Ala and Gly insertions have different predicted ABEGO classifications, ColabFold primarily models the Gly conformation to be in the G region (10/11 sites in different ABEGO bins where the Gly conformation is G). This suggests that our observed differences between Ala and Gly insertions owe to Glycine’s ability to adopt a greater range of backbone conformations.

**Figure 6.**
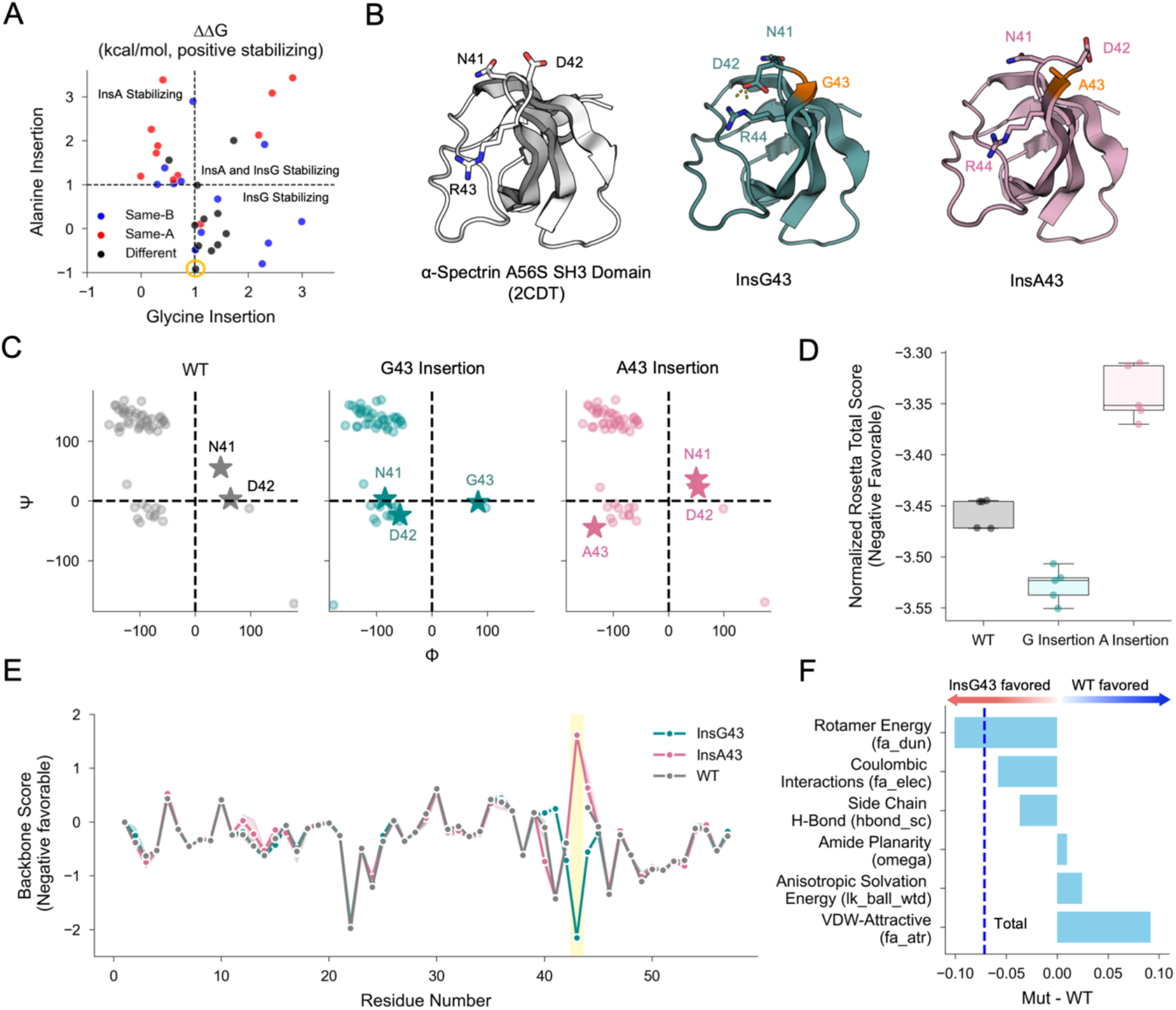
**A**, Relationship between ΔΔG_unfolding_ of alanine and glycine insertions for 36 sites stabilized by at least one insertion. Points are colored by ABEGO agreement between the alanine and glycine insertions in the ColabFold modeled structures. Yellow circle highlights alanine and glycine insertions at position 43 in A56S α-spectrin (2CDT). **B**, ColabFold models of A56S α-spectrin (2CDT) and insertion mutants InsG43 and InsA43. **C**, Ramachandran maps of 2CDT and insertion mutants InsG43 and InsA43. Residues N41, D42, A43, and G43 are represented as stars. **D**, Mean normalized Rosetta total scores for the five lowest-scoring relaxed structures of 2CDT, InsG43, and InsA43. **E**, Per-residue backbone score profiles for 2CDT, InsG43, and InsA43. Residue 43 is highlighted in yellow. **F**, Normalized changes in Rosetta score terms between insertion mutant InsG43 and wild-type 2CDT. The normalized change in Rosetta total score between 2CDT and InsG43 is shown by the vertical dashed blue line.

The α-spectrin SH3 domain A56S mutant (PDB ID: 2CDT) illustrates how Gly insertions can stabilize a domain when Ala insertions are not tolerated. α-spectrin is a scaffold protein involved in crosslinking the plasma membrane to the actin cytoskeleton and is comprised of multiple modular protein domains, including a highly conserved SRC Homology 3 (SH3) domain (*51*). The A56S mutant of this domain was previously studied to understand the folding stability of the domain (*52*). We refer to this mutant as the “wild type” because this sequence was used as the background for the deep mutational scan in the Tsuboyama et al. dataset (Fig. 6B). This domain is stabilized by the InsG43 mutation (ΔΔG_Gly_ = 1.0 kcal/mol), but is destabilized by an Alanine insertion in the same position (ΔΔG_Ala_ = -1.2 kcal/mol) (Fig 6A, yellow circle). The stabilizing effect of the InsG43 mutant may owe to that mutant’s ability to relieve the strain on N41 and D42. In the wild-type sequence, N41 and D42 are located in the positive phi (G) region of the Ramachandran map, which is unfavorable (Fig. 6C). The Gly43 insertion (but not the Ala43 insertion) is predicted to shift these residues to the more favorable negative phi region. This is made possible by the inserted Gly43 adopting a positive phi angle (G region), which is not predicted for an inserted Ala43 residue.

Although these predicted changes in backbone geometry may explain the differing effects of InsG43 and InsA43, Rosetta’s physics-based model offers a different explanation. Rosetta correctly predicts the InsG43 mutant to be more favorable (by total score) than the wild type, and the InsA43 mutant to be less favorable than the wild type (Fig. 6D). In analyzing the per-residue backbone scores for InsG43, InsA43, and the wild type, we see that Rosetta strongly prefers an inserted glycine in position 43 compared to alanine (Fig. 6E). However, N41 becomes more unfavorable in the InsG43 mutant compared to the wild type and InsA43 mutant in backbone score (Fig. 6E). Despite this, Rosetta favors InsG43 due to a combination of more favorable side chain conformations and interactions, instead of a change in backbone strain (Fig. 6F). This highlights the complexity of interpreting and predicting changes in protein energetics resulting from indel mutations. While some indels likely increase folding stability by relieving backbone strain (as suggested in the Pin1 WW domain), the case of the α-spectrin SH3 domain illustrates that the energetic effects of stabilizing indels cannot solely be explained by backbone strain.

## Conclusions

We analyzed structural and energetic changes for 103 stabilizing insertion and deletion mutations to understand how indels can increase stability and reveal the role of backbone energetics on global stability. We found that stabilizing indel mutations tend to have minimal structural perturbations and are frequently found in loops, but can also be found in secondary structure elements as well. Stabilizing indels are typically correctly classified as stabilizing by the Rosetta energy function, but not by the Cagiada et al. model based on ESM-IF (*35*). Unlike ESM-IF, Rosetta explicitly calculates this backbone strain, which may explain why Rosetta is more successful at classifying stabilizing indel variants. Using the Rosetta energy function, we discovered that stabilizing insertions and deletions are often found in regions of high backbone strain. Finally, we examined two case studies of stabilizing indel mutations: the Pin1 WW domain and the α-spectrin SH3 domain.

Our analysis has several limitations. First, the structures of our indel mutants and wild-type domains were predicted using ColabFold, which is informed by evolutionary information of the sequence. These predictions may not correctly model sequence variants or proteins with little to no evolutionary information. Second, we only evaluated Rosetta and the ESM-IF-based model at predicting the mutational effects of stabilizing indels. Whereas we found that Rosetta had fewer “false negatives” (i.e. predicting stabilizing indels as destabilizing), both Rosetta and ESM-IF may predict “false positives” (predicting destabilizing indels as stabilizing). Third, our backbone strain calculations are based on static models that we assume represent energy minima, and our analysis does not consider the full conformational ensembles of these domains. Finally, although we primarily focus on backbone strain, indels affect protein energetics in complex ways, as illustrated by the α-spectrin SH3 domain. In particular, indels of varying lengths and insertions of amino acids other than alanine or glycine may affect protein energetics differently from our results.

Indels can improve a protein’s folding stability by favorably changing its backbone geometry. We find that Rosetta more accurately predicts the mutational effects of stabilizing indels than the ESM-IF analysis. However, Rosetta only correctly classified approximately two-thirds of stabilizing mutants, highlighting the need for more accurate models in predicting ΔG and ΔΔG. Because stabilizing indels are rare and necessitate changes to backbone geometry, these mutations serve as challenging test cases for prediction models. These mutations reveal the limitations of current methods and can be used to benchmark future methods. By collecting more data about the mutational effects of indels, and developing better predictive models, we can better leverage these mutations in protein design efforts.

## Acknowledgments

We thank Állan Ferrari for helpful discussions and technical support using LovoAlign and Rosetta. We are grateful to members of the Rocklin Lab for their comments and suggestions on this manuscript. This research was supported in part through the computational resources and staff contributions provided for the Quest high performance computing facility at Northwestern University which is jointly supported by the Office of the Provost, the Office for Research, and Northwestern University Information Technology. Y.M.G was supported by the Chemistry of Life Processes CAURS Undergraduate Research Award and is currently supported by the Shurl and Kay Curci Foundation PhD Scholarship. G.J.R. acknowledges support from the National Institutes of Health under award number DP2GM140927.

## Methods

### Dataset

The Mega-Scale dataset from Tsuboyama et al., 2023 was downloaded from the April 2023 updated release on Zenodo (https://zenodo.org/records/7992926). We used Dataset 3, which reports 607,839 stability measurements of 412 wild-type domains. Unreliable ΔG entries, marked with ‘-’, were removed, resulting in a dataset of 57,698 high-quality measurements of single amino acid insertion and deletion mutants (indels) of 405 wild types. Seven domains had no indel mutants with reliable ΔG entries. From this filtered dataset of indels, 108 stabilizing mutants (ΔΔG > 1 kcal/mol) were selected. Five stabilizing deletion mutants (EHEE_rd1_0101_delR38, EHEE_rd1_0101_delR39, EHEE_rd1_0407_delD40, 1SF0_V59K_delK59, and 1SF0_V59K_delP63) were removed because nearly any substitution or indel mutation at these sites was stabilizing, suggesting the improved stability was an artifact of changing a protease cleavage site. This filtering yielded a final dataset of 103 stabilizing indel mutants of 48 wild types.

### Structure Prediction

Mutant and wild-type structures were predicted using ColabFold. We ran ColabFold using default parameters and selected the top-ranking (lowest average pLDDT) model for analysis. We did not generate multiple sequence alignments (MSAs) for designed sequences. Stabilizing mutant and wild-type structures with an average pLDDT < 85 and RMSD > 5 Å were excluded from structural and energetic analyses, yielding a final set of 92 indel mutants for structural and energetic analyses.

### Structural Analysis

The secondary structure of inserted and deleted residues was determined using the Dssp module in PyRosetta.

Stabilizing indel mutants and corresponding wild-type predicted structures were aligned using LovoAlign (https://m3g.github.io/lovoalign/).

### Energetic Analysis

#### Stability Predictions using the Cagiada ESM-IF Analysis

92 indel and wild-type ColabFold models were evaluated using the Cagiada ESM-IF analysis (https://github.com/KULL-Centre/_2024_cagiada_stability/). Absolute stability predictions (ΔG_unfolding_) were made using ESM-IF for the indel mutants and wild types. Stability predictions were normalized by protein length to account for the length dependence of the Cagiada ESM-IF analysis.

#### Rosetta Scoring

Predicted structures were relaxed and scored using the Rosetta Fast Relax protocol. 20 relax runs were performed for each predicted structure. Scores for each predicted structure were calculated by averaging individual score terms from the five lowest-scoring (by total energy) relaxed structures. Rosetta protein-level scores were normalized by dividing by protein length.

## References

1. R. Kim, J.-T. Guo, Systematic analysis of short internal indels and their impact on protein folding. BMC Struct. Biol. 10, 1–11 (2010).

2. M. Lin, S. Whitmire, J. Chen, A. Farrel, X. Shi, J.-T. Guo, Effects of short indels on protein structure and function in human genomes. Sci. Rep. 7, 1–9 (2017).

3. S. Savino, T. Desmet, J. Franceus, Insertions and deletions in protein evolution and engineering. Biotechnol. Adv. 60, 108010 (2022).

4. B. Danneels, M. Pinto-Carbó, A. Carlier, Patterns of nucleotide deletion and insertion inferred from bacterial pseudogenes. Genome Biol. Evol. 10, 1792–1802 (2018).

5. M. Lynch, M. S. Ackerman, J.-F. Gout, H. Long, W. Sung, W. K. Thomas, P. L. Foster, Genetic drift, selection and the evolution of the mutation rate. Nat. Rev. Genet. 17, 704– 714 (2016).

6. G. L. Lukacs, A. S. Verkman, CFTR: folding, misfolding and correcting the ΔF508 conformational defect. Trends Mol. Med. 18, 81–91 (2012).

7. K. Nishitsuji, T. Tomiyama, K. Ishibashi, K. Ito, R. Teraoka, M. P. Lambert, W. L. Klein, H. Mori, The E693Delta mutation in amyloid precursor protein increases intracellular accumulation of amyloid beta oligomers and causes endoplasmic reticulum stress-induced apoptosis in cultured cells. Am. J. Pathol. 174, 957–969 (2009).

8. M. Seuma, B. Lehner, B. Bolognesi, An atlas of amyloid aggregation: the impact of substitutions, insertions, deletions and truncations on amyloid beta fibril nucleation. Nat. Commun. 13, 7084 (2022).

9. M. Chansel-Da Cruz, M. Hohl, I. Ceppi, L. Kermasson, L. Maggiorella, M. Modesti, J.-P. de Villartay, T. Ileri, P. Cejka, J. H. J. Petrini, P. Revy, A Disease-Causing Single Amino Acid Deletion in the Coiled-Coil Domain of RAD50 Impairs MRE11 Complex Functions in Yeast and Humans. Cell Rep. 33, 108559 (2020).

10. M. Topolska, A. Beltran, B. Lehner, Deep indel mutagenesis reveals the impact of amino acid insertions and deletions on protein stability and function, bioRxiv, doi: 10.1101/2023.10.06.561180 (2024).

11. C. M. Miton, N. Tokuriki, Insertions and deletions (indels): A missing piece of the protein engineering jigsaw. Biochemistry 62, 148–157 (2023).

12. D. Shortle, J. Sondek, The emerging role of insertions and deletions in protein engineering. Curr. Opin. Biotechnol. 6, 387–393 (1995).

13. J. L. Tenthorey, S. Del Banco, I. Ramzan, H. Klingenberg, C. Liu, M. Emerman, H. S. Malik, Indels allow antiviral proteins to evolve functional novelty inaccessible by missense mutations, bioRxiv, doi: 10.1101/2024.05.07.592993 (2024).

14. S. Emond, M. Petek, E. J. Kay, B. Heames, S. R. A. Devenish, N. Tokuriki, F. Hollfelder, Accessing unexplored regions of sequence space in directed enzyme evolution via insertion/deletion mutagenesis. Nat. Commun. 11, 3469 (2020).

15. G. S. Baird, D. A. Zacharias, R. Y. Tsien, Circular permutation and receptor insertion within green fluorescent proteins. Proc. Natl. Acad. Sci. U. S. A. 96, 11241–11246 (1999).

16. J. A. J. Arpino, S. C. Reddington, L. M. Halliwell, P. J. Rizkallah, D. D. Jones, Random single amino acid deletion sampling unveils structural tolerance and the benefits of helical registry shift on GFP folding and structure. Structure 22, 889–898 (2014).

17. T. Grocholski, P. Dinis, L. Niiranen, J. Niemi, M. Metsä-Ketelä, Divergent evolution of an atypical S-adenosyl-l-methionine-dependent monooxygenase involved in anthracycline biosynthesis. Proc. Natl. Acad. Sci. U. S. A. 112, 9866–9871 (2015).

18. D. M. Fowler, S. Fields, Deep mutational scanning: a new style of protein science. Nat. Methods 11, 801–807 (2014).

19. M. Chiasson, M. J. Dunham, A. E. Rettie, D. M. Fowler, Applying multiplex assays to understand variation in pharmacogenes. Clin. Pharmacol. Ther. 106, 290–294 (2019).

20. C. B. Macdonald, D. Nedrud, P. R. Grimes, D. Trinidad, J. S. Fraser, W. Coyote-Maestas, DIMPLE: deep insertion, deletion, and missense mutation libraries for exploring protein variation in evolution, disease, and biology. Genome Biol. 24, 36 (2023).

21. C. E. Gonzalez, P. Roberts, M. Ostermeier, Fitness effects of single amino acid insertions and deletions in TEM-1 β-lactamase. J. Mol. Biol. 431, 2320–2330 (2019).

22. K. Skamaki, S. Emond, M. Chodorge, J. Andrews, D. G. Rees, D. Cannon, B. Popovic, A. Buchanan, R. R. Minter, F. Hollfelder, In vitro evolution of antibody affinity via insertional scanning mutagenesis of an entire antibody variable region. Proc. Natl. Acad. Sci. U. S. A. 117, 27307–27318 (2020).

23. A. Schenkmayerova, G. P. Pinto, M. Toul, M. Marek, L. Hernychova, J. Planas-Iglesias, V. Daniel Liskova, D. Pluskal, M. Vasina, S. Emond, M. Dörr, R. Chaloupkova, D. Bednar, Z. Prokop, F. Hollfelder, U. T. Bornscheuer, J. Damborsky, Engineering the protein dynamics of an ancestral luciferase. Nat. Commun. 12, 3616 (2021)

24. L. Rockah-Shmuel, Á. Tóth-Petróczy, A. Sela, O. Wurtzel, R. Sorek, D. S. Tawfik, Correlated occurrence and bypass of frame-shifting insertion-deletions (InDels) to give functional proteins. PLoS Genet. 9, e1003882 (2013).

25. C. Douville, D. L. Masica, P. D. Stenson, D. N. Cooper, D. M. Gygax, R. Kim, M. Ryan, R. Karchin, Assessing the pathogenicity of insertion and deletion variants with the variant effect scoring tool (VEST-indel). Hum. Mutat. 37, 28–35 (2016).

26. A. M. Simm, A. J. Baldwin, K. Busse, D. D. Jones, Investigating protein structural plasticity by surveying the consequence of an amino acid deletion from TEM-1 beta-lactamase. FEBS Lett. 581, 3904–3908 (2007).

27. L. M. Halliwell, A. P. Jathoul, J. P. Bate, H. L. Worthy, J. C. Anderson, D. D. Jones, J. A. H. Murray, ΔFlucs: Brighter Photinus pyralis firefly luciferases identified by surveying consecutive single amino acid deletion mutations in a thermostable variant. Biotechnol. Bioeng. 115, 50–59 (2018).

28. S. Larsen-Ledet, S. Lindemose, A. Panfilova, S. Gersing, C. H. Suhr, A. V. Genzor, H. Lanters, S. V. Nielsen, K. Lindorff-Larsen, J. R. Winther, A. Stein, R. Hartmann-Petersen, Systematic characterization of indel variants using a yeast-based protein folding sensor, bioRxiv, doi: 10.1101/2024.07.11.603017 (2024).

29. P. J. Ogden, E. D. Kelsic, S. Sinai, G. M. Church, Comprehensive AAV capsid fitness landscape reveals a viral gene and enables machine-guided design. Science 366, 1139– 1143 (2019).

30. S. Pascarella, P. Argos, Analysis of insertions/deletions in protein structures. J. Mol. Biol. 224, 461–471 (1992).

31. J. Hu, P. C. Ng, SIFT Indel: predictions for the functional effects of amino acid insertions/deletions in proteins. PLoS One 8, e77940 (2013).

32. Y. Choi, G. E. Sims, S. Murphy, J. R. Miller, A. P. Chan, Predicting the functional effect of amino acid substitutions and indels. PLoS One 7, e46688 (2012).

33. S. Dagan, T. Hagai, Y. Gavrilov, R. Kapon, Y. Levy, Z. Reich, Stabilization of a protein conferred by an increase in folded state entropy. Proc. Natl. Acad. Sci. U. S. A. 110, 10628– 10633 (2013).

34. K. Tsuboyama, J. Dauparas, J. Chen, E. Laine, Y. Mohseni Behbahani, J. J. Weinstein, N. M. Mangan, S. Ovchinnikov, G. J. Rocklin, Mega-scale experimental analysis of protein folding stability in biology and design. Nature 620, 434–444 (2023).

35. M. Cagiada, S. Ovchinnikov, K. Lindorff-Larsen, Predicting absolute protein folding stability using generative models, bioRxiv, doi: 10.1101/2024.03.14.584940 (2024).

36. A. Federico, R. Sepe, F. Cozzolino, C. Piccolo, C. Iannone, I. Iacobucci, P. Pucci, M. Monti, A. Fusco, The complex CBX7-PRMT1 has a critical role in regulating E-cadherin gene expression and cell migration. Biochim. Biophys. Acta Gene Regul. Mech. 1862, 509–521 (2019).

37. M. Mirdita, K. Schütze, Y. Moriwaki, L. Heo, S. Ovchinnikov, M. Steinegger, ColabFold: making protein folding accessible to all. Nat. Methods 19, 679–682 (2022).

38. L. Martínez, R. Andreani, J. M. Martínez, Convergent algorithms for protein structural alignment. BMC Bioinformatics 8, 306 (2007).

39. J. K. Leman, B. D. Weitzner, S. M. Lewis, J. Adolf-Bryfogle, N. Alam, R. F. Alford, M. Aprahamian, D. Baker, K. A. Barlow, P. Barth, B. Basanta, B. J. Bender, K. Blacklock, J. Bonet, S. E. Boyken, P. Bradley, C. Bystroff, P. Conway, S. Cooper, B. E. Correia, B. Coventry, R. Das, R. M. De Jong, F. DiMaio, L. Dsilva, R. Dunbrack, A. S. Ford, B. Frenz, D. Y. Fu, C. Geniesse, L. Goldschmidt, R. Gowthaman, J. J. Gray, D. Gront, S. Guffy, S. Horowitz, P.-S. Huang, T. Huber, T. M. Jacobs, J. R. Jeliazkov, D. K. Johnson, K. Kappel, J. Karanicolas, H. Khakzad, K. R. Khar, S. D. Khare, F. Khatib, A. Khramushin, I. C. King, R. Kleffner, B. Koepnick, T. Kortemme, G. Kuenze, B. Kuhlman, D. Kuroda, J. W. Labonte, J. K. Lai, G. Lapidoth, A. Leaver-Fay, S. Lindert, T. Linsky, N. London, J. H. Lubin, S. Lyskov, J. Maguire, L. Malmström, E. Marcos, O. Marcu, N. A. Marze, J. Meiler, R. Moretti, V. K. Mulligan, S. Nerli, C. Norn, S. ó’Conchúir, N. Ollikainen, S. Ovchinnikov, M. S. Pacella, X. Pan, H. Park, R. E. Pavlovicz, M. Pethe, B. G. Pierce, K. B. Pilla, B. Raveh, P. D. Renfrew, S. S. R. Burman, A. Rubenstein, M. F. Sauer, A. Scheck, W. Schief, O. Schueler-Furman, Y. Sedan, A. M. Sevy, N. G. Sgourakis, L. Shi, J. B. Siegel, D.-A. Silva, S. Smith, Y. Song, A. Stein, M. Szegedy, F. D. Teets, S. B. Thyme, R. Y.-R. Wang, A. Watkins, L. Zimmerman, R. Bonneau, Macromolecular modeling and design in Rosetta: recent methods and frameworks. Nat. Methods 17, 665–680 (2020).

40. R. F. Alford, A. Leaver-Fay, J. R. Jeliazkov, M. J. O’Meara, F. P. DiMaio, H. Park, M. V. Shapovalov, P. D. Renfrew, V. K. Mulligan, K. Kappel, J. W. Labonte, M. S. Pacella, R. Bonneau, P. Bradley, R. L. Dunbrack Jr, R. Das, D. Baker, B. Kuhlman, T. Kortemme, J. J. Gray, The Rosetta all-atom energy function for macromolecular modeling and design. J. Chem. Theory Comput. 13, 3031–3048 (2017).

41. Y. Chen, Y.-R. Wu, H.-Y. Yang, X.-Z. Li, M.-M. Jie, C.-J. Hu, Y.-Y. Wu, S.-M. Yang, Y.-B. Yang, Prolyl isomerase Pin1: a promoter of cancer and a target for therapy. Cell Death Dis. 9, 883 (2018).

42. M. Jäger, Y. Zhang, J. Bieschke, H. Nguyen, M. Dendle, M. E. Bowman, J. P. Noel, M. Gruebele, J. W. Kelly, Structure-function-folding relationship in a WW domain. Proc. Natl. Acad. Sci. U. S. A. 103, 10648–10653 (2006).

43. M. Jäger, M. Dendle, J. W. Kelly, Sequence determinants of thermodynamic stability in a WW domain--an all-beta-sheet protein. Protein Sci. 18, 1806–1813 (2009).

44. M. Jäger, H. Nguyen, J. C. Crane, J. W. Kelly, M. Gruebele, The folding mechanism of a beta-sheet: the WW domain. J. Mol. Biol. 311, 373–393 (2001).

45. Y. M. Lee, Y.-C. Liou, Gears-in-motion: The interplay of WW and PPIase domains in Pin1. Front. Oncol. 8, 469 (2018).

46. C. L. Araya, D. M. Fowler, W. Chen, I. Muniez, J. W. Kelly, S. Fields, A fundamental protein property, thermodynamic stability, revealed solely from large-scale measurements of protein function. Proc. Natl. Acad. Sci. U. S. A. 109, 16858–16863 (2012).

47. R. Wintjens, J. M. Wieruszeski, H. Drobecq, P. Rousselot-Pailley, L. Buée, G. Lippens, I. Landrieu, 1H NMR study on the binding of Pin1 Trp-Trp domain with phosphothreonine peptides. J. Biol. Chem. 276, 25150–25156 (2001).

48. M. A. Verdecia, M. E. Bowman, K. P. Lu, T. Hunter, J. P. Noel, Structural basis for phosphoserine-proline recognition by group IV WW domains. Nat. Struct. Biol. 7, 639– 643 (2000).

49. T. Peng, J. S. Zintsmaster, A. T. Namanja, J. W. Peng, Sequence-specific dynamics modulate recognition specificity in WW domains. Nat. Struct. Mol. Biol. 14, 325–331 (2007).

50. Y.-R. Lin, N. Koga, R. Tatsumi-Koga, G. Liu, A. F. Clouser, G. T. Montelione, D. Baker, Control over overall shape and size in de novo designed proteins. Proc. Natl. Acad. Sci. U. S. A. 112, E5478–85 (2015).

51. B. Machnicka, R. Grochowalska, D. M. Bogusławska, A. F. Sikorski, The role of spectrin in cell adhesion and cell-cell contact. Exp. Biol. Med. 244, 1303–1312 (2019).

52. S. Casares, O. López-Mayorga, M. C. Vega, A. Cámara-Artigas, F. Conejero-Lara, Cooperative propagation of local stability changes from low-stability and high-stability regions in a SH3 domain. Proteins 67, 531–547 (2007).

